# Characterizing the effects of sex, *APOE* ɛ4, and literacy on mid-life cognitive trajectories: Application of Information-Theoretic model-averaging and multi-model inference techniques to the Wisconsin Registry for Alzheimer’s Prevention Study

**DOI:** 10.1101/229237

**Authors:** Rebecca L. Koscik, Derek L. Norton, Samantha L. Allison, Erin M. Jonaitis, Lindsay R. Clark, Kimberly D. Mueller, Bruce P. Hermann, Corinne D. Engelman, Carey E. Gleason, Mark A. Sager, Richard J. Chappell, Sterling C. Johnson

**Affiliations:** Wisconsin Alzheimer’s Institute, University of Wisconsin School of Medicine and Public Health, Madison, WI, USA; Department of Biostatistics and Medical Informatics, University of Wisconsin School of Medicine and Public Health, Madison, WI USA; Geriatric Research Education and Clinical Center, William S. Middleton Memorial Veterans Hospital, Madison WI, USA; Alzheimer’s Disease Research Center, University of Wisconsin-Madison School of Medicine and Public Health, Madison, WI, USA; Department of Neurology, University of Wisconsin School of Medicine and Public Health, Madison, WI 53705, USA; Department of Population Health Sciences, University of Wisconsin School of Medicine and Public Health, Madison, WI 53726, USA; Department of Statistics, University of Wisconsin-Madison, Madison, WI 53706, USA

**Author notes:** **Corresponding author:** Rebecca Koscik, 610 Walnut St. (Rm. 944), Madison, WI, 53726 Office.

**Keywords:** Kullback-Leibler divergence, model averaging, model likelihoods, model selection, cognitive decline, Alzheimer’s disease

## Abstract

**Objective:** In this paper we apply Information-Theoretic (IT) model averaging to characterize a set of complex interactions in a longitudinal study on cognitive decline. Prior research has identified numerous genetic (including sex), education, health and lifestyle factors that predict cognitive decline. Traditional model selection approaches (e.g., backward or stepwise selection) attempt to find models that best fit the observed data; these techniques risk interpretations that only the selected predictors are important. In reality, several models may fit similarly well but result in different conclusions (e.g., about size and significance of parameter estimates); inference from traditional model selection approaches can lead to overly confident conclusions.

**Method:** Here we use longitudinal cognitive data from ~1550 late-middle aged adults the Wisconsin Registry for Alzheimer’s Prevention study to examine the effects of sex, Apolipoprotein E (*APOE*) ɛ4 allele (non-modifiable factors), and literacy achievement (modifiable) on cognitive decline. For each outcome, we applied IT model averaging to a model set with combinations of interactions among sex, *APOE*, literacy, and age.

**Results:** For a list-learning test, model-averaged results showed better performance for women vs men, with faster decline among men; increased literacy was associated with better performance, particularly among men. *APOE* had less of an effect on cognitive performance in this age range (~40-70).

**Conclusions:** These results illustrate the utility of the IT approach and point to literacy as a potential modifier of decline. Whether the protective effect of literacy is due to educational attainment or intrinsic verbal intellectual ability is the topic of ongoing work.

## Introduction

Signs of cognitive decline often begin a decade or more before diagnosis of dementia due to late-onset Alzheimer’s disease (AD), a neurodegenerative disease associated with greatly impaired cognition and daily functioning (Price et al., 2009; Price & Morris, 1999). After age, presence of one or more Apolipoprotein E (*APOE*) ɛ4 alleles is the strongest predictor of risk of late onset AD (Strittmatter & Roses, 1996; Tang et al., 1996), particularly among non-Hispanic Caucasians (Tang et al., 1998). Several studies report earlier and faster declines in memory or executive function among *APOE* ɛ4 carriers than non-carriers (i.e., *APOE* by age interactions), with detectable accelerations in decline beginning around age 60 (Caselli et al., 2009; Chang et al., 2014; Wisdom, Callahan, & Hawkins, 2011). Studies also indicate that the rate of ɛ4-associated cognitive decline and AD risk are moderated by sex with ɛ4-carriage increasing rate of decline and risk more in women than in men (Altmann, Tian, Henderson, & Greicius, 2014; Beydoun et al., 2012; Koran, Wagener, & Hohman, 2017; Mielke, Vemuri, & Rocca, 2014; Mortensen & Høgh, 2001; Neu et al., 2017; Payami et al., 1996; Riedel, Thompson, & Brinton, 2016). Higher literacy levels (as measured by word reading tasks) have been shown to mitigate age - and/or *APOE*-related cognitive decline in non-demented elders (Kaup et al., 2015; Manly, Touradji, Tang, & Stern, 2003).

No studies, to our knowledge, have investigated the *combined* influences of sex, *APOE* genotype, and literacy on early to late middle-age cognitive trajectories in an integrated analytic framework. Traditional model selection approaches, such as forward, backward, or stepwise selection, attempt to find a model that best fits the observed data. One issue that can arise from such methods is concluding that only the selected predictors are important while assuming those not selected are unimportant (D. R. Anderson & Burnham, 2002). In reality, several models may fit the data similarly well but result in different conclusions, and inference from a single model chosen after a selection procedure can lead to overly optimistic results and conclusions (Claeskens and Hjort 2008). Information-theoretic (IT) modeling techniques offer a way to characterize complex sets of interactions and make multi-model inference while avoiding the pitfalls of predictor selection methods (D. R. Anderson & Burnham, 2002; Claeskens & Hjort, 2008).

The IT framework focus is on “a small set of science hypotheses, all of which are plausible” (p. 202, (D. R. Anderson & Burnham, 2002)). The IT approach has its roots in biological ecology research (Kenneth P. Burnham, Anderson, & Huyvaert, 2011; Hegyi & Garamszegi, 2011; Richards, 2005; Richards, Whittingham, & Stephens, 2011; Symonds & Moussalli, 2011), and aims to use the relative strength of information among all considered models instead of selecting a single model. The methods start with formulating a reasonably sized collection of models with the same outcome, but different covariate structures (such as different main effects, interactions, etc.). The differences between the models should all be items of interest and scientifically reasonable to include. After fitting all of these models to the data, results are combined across models in proportion to the relative strength of information each model provides. These relative strengths are quantified through the theory of the corrected Akaike Information Criterion (AICc), which estimates the relative differences among these models with respect to their Kullback-Leibler divergence, a measure of the distance between a proposed model and the “true” model (Hurvich & Tsai, 1989). This allows models fitting similarly well to contribute relatively equal amounts of influence on the resulting parameter estimates, while models that fit poorly have little or no influence on results.

The aim of this study was to use IT model-averaging to characterize how sex, literacy, and *APOE* genotype influence age-related trajectories for several neuropsychological tests in a longitudinal sample enriched for risk of developing AD (Wisconsin Registry for Alzheimer’s Prevention (WRAP)). In secondary analyses, we compare the IT model-averaging results with traditional model selection methods.

## Methods

### WRAP study and participants

WRAP is a longitudinal cohort study enriched for AD-risk via over-enrollment of participants with a parental history of AD (for details, see (Johnson et al., 2017)); primary aims of the study include identifying predictors associated with cognitive decline and estimating their associations. All participants were free of dementia at baseline. At the time of these analyses, there were 1549 enrolled WRAP participants (baseline age mean(sd)= 53.7(6.6); parental history of AD n(%)= 1125(72.6%)). To be included in these analyses, participants had to be free at baseline of MCI and any of four neurological conditions (stroke, Parkinson’s disease, multiple sclerosis, epilepsy), have completed at least 2 study visits, and have complete data in the predictors needed for the analyses (n=1256 eligible; n’s excluded: MCI, n=4; neurological disorder, n=59; <2 visits, n=226; incomplete predictors, n=4). Depending upon the outcome, additional subjects were excluded due to missing outcome values resulting in <2 visits with a value (details in Table 1).

**Table 1:** Sample characteristics

|  | Overall | Female | Male | p-value* |
| --- | --- | --- | --- | --- |
| N | 1256 | 873 (69.5%) | 383 (30.5%) |  |
| Age (mean (sd)) | 53.67 (6.55) | 53.56 (6.57) | 53.92 (6.51) | 0.238 |
| URG (N (%)) | 85 (6.8) | 61 (7.0) | 24 (6.3) | 0.715+ |
| ESL (N (%)) | 21 (1.7) | 16 (1.8) | 5 (1.3) | 0.636+ |
| Follow-up years (mean (sd)) | 9.10 (2.60) | 9.04 (2.63) | 9.22 (2.53) | 0.34 |
| Number of visits (N (%)) |  |  |  |  |
| 2 | 135 (10.7) | 102 (11.7) | 33 (8.6) | 0.124+ |
| 3 | 265 (21.1) | 180 (20.6) | 85 (22.2) |  |
| 4 | 432 (34.4) | 310 (35.5) | 122 (31.9) |  |
| 5 | 424 (33.8) | 281 (32.2) | 143 (37.3) |  |
| APOE ε4 count (N (%)) |  |  |  |  |
| 0 | 767 (61.1) | 523 (59.9) | 244 (63.7) | <0.001+ |
| 1 | 438 (34.9) | 303 (34.7) | 135 (35.2) |  |
| 2 | 51 (4.1) | 47 (5.4) | 4 (1.0) |  |
| SES (N (%)) |  |  |  |  |
| 1 | 39 (3.1) | 26 (3.0) | 13 (3.4) | <0.001^ |
| 2 | 122 (9.7) | 99 (11.3) | 23 (6.0) |  |
| 3 | 267 (21.3) | 202 (23.1) | 65 (17.0) |  |
| 4 | 241 (19.2) | 171 (19.6) | 70 (18.3) |  |
| 5 | 587 (46.7) | 375 (43.0) | 212 (55.4) |  |
| RAVLT total (mean (sd)) | 51.10 (7.95) | 52.61 (7.30) | 47.64 (8.30) | <0.001 |
| RAVLT delayed recall (mean (sd)) | 10.46 (2.82) | 10.96 (2.60) | 9.33 (2.99) | <0.001 |
| Trails A (mean (sd)) | 26.81 (8.55) | 26.28 (8.24) | 28.02 (9.11) | 0.001 |
| Trails B (mean (sd)) | 62.06 (24.61) | 60.21 (22.16) | 66.28 (29.03) | <0.001 |
| Boston Naming 60 (mean (sd)) | 57.02 (3.09) | 56.87 (3.19) | 57.35 (2.84) | 0.002 |
| CFL fluency (mean (sd)) | 43.37 (11.07) | 43.99 (10.75) | 41.94 (11.65) | 0.002 |
| Digit Span backward (mean (sd)) | 7.09 (2.22) | 7.00 (2.17) | 7.31 (2.32) | 0.024 |
| Digit Span forward (mean (sd)) | 10.51 (2.18) | 10.40 (2.11) | 10.76 (2.31) | 0.006 |
| Stroop color word, (mean (sd)) | 108.12 (20.63) | 109.86 (19.58) | 104.09 (22.42) | <0.001 |
| 9-Test IICV (mean (sd)) | 0.81 (0.23) | 0.80 (0.22) | 0.82 (0.24) | 0.069 |
| 4-Test IICV (mean (sd)) | 0.73 (0.33) | 0.73 (0.33) | 0.73 (0.33) | 0.803 |

\*Note: Comparisons made with Mann-Whitney, unless noted with + (Fisher's exact) or ^ (Chi-squared);

### Study protocol and outcomes

At each study visit, participants completed comprehensive cognitive assessments, detailed health and lifestyle questionnaires, and provided blood samples for current and future analyses. The first follow-up visit occurred approximately 4 years after baseline, with subsequent visits occurring approximately every 2 years (for details, see (Johnson et al., 2017)). These analyses focus on nine cognitive measures available since baseline of the WRAP study and shown to be sensitive to AD-related cognitive changes. The tests and measures used are: the Rey Auditory Verbal Learning and Memory Test (Schmidt, 1996), sum of learning trials (“AVLT Total”) and long delay recall trial (“AVLT Delay”); Trail Making Test (“Trails A” and “Trails B”, (Lezak, Howieson, Bigler, & Tranel, 2012)); Stroop Color-Word Interference Test (Trenerry, Crosson, DeBoe, & Leber, 1989), number of correct items in two minutes; Controlled Oral Word Association Test (Benton, Hamsher, & Sivan, 1994), total words in 60 seconds for each letter: C, F, L (“CFL”); the Boston Naming Test (“BNT”, (Kaplan, Goodglass, & Weintraub, 2001)), total correct and Digit span forward and backward total items correct (Wechsler, 1997).

Given recent results in WRAP and other studies suggesting that higher intraindividual cognitive variability (IICV) at a given visit predicts increased risk of subsequent decline (E. D. Anderson et al., 2016; Gleason et al., 2017; Holtzer, Verghese, Wang, Hall, & Lipton, 2008; Koscik et al., 2016), we also characterized how IICV varied by sex, literacy, *APOE*, and age in our sample. We calculated two versions of IICV. The first represents the version we used in a previous paper (Koscik et al., 2016), “4-Test IICV”, and was calculated as the standard deviation of z-scores of AVLT Total, Trails A and B, and the Wide Range Achievement Test (3^rd^ ed., “WRAT”) reading subtest standard score (Wilkinson, 1993). The WRAT Reading Achievement score when used in middle-aged and older adults is accepted as a proxy for maximal verbal intellectual attainment and quality of education. Higher WRAT Reading scores have been associated with slower memory declines in non-demented elders (Manly et al. 2003). The second IICV version was calculated as the standard deviation of z-scores of the nine tests selected for analysis in this manuscript (9-Test IICV). Before z-scoring occurred, all tests were assessed for normality, and Box-Cox transformed for non-normal items (Trails A and B, BNT, Digit Span Forward, WRAT, and AVLT Delay+1).

### Key Predictors and Covariates

Key predictors in this study include age (years), chromosomal sex (Male/Female), *APOE* ɛ4 allele count (i.e., 0, 1, 2 (for details on genotyping methods, see (Darst et al., 2017))), and WRAT. For subjects without baseline WRAT, the value at their second visit was used. Additional covariates included race/ethnicity (non-Hispanic Caucasian vs underrepresented group, URG), if English was the subject’s native language (ESL), and socioeconomic status (SES; : 1 = <$20k; 2 = $20k-<$40k; 3= $40k-<$60k; 4=$60k-<$80k, and 5= $80k or more). Missingness in SES was present in 54 (4.3%) participants. To recover these subjects for analyses, their baseline SES values were imputed through proportional odds regression using baseline values of age, chromosomal sex, race/ethnicity, the Center for Epidemiological Studies of Depression (CES-D) total score (Radloff, 1977), literacy, and years of education as predictors.

### Data analysis

We followed the steps outlined for the IT-modeling approach detailed below.

#### 1. Specifying the model set

Based on research indicating potential interactions between literacy and sex, *APOE* ɛ4 count or age (on cognitive outcomes), we developed a set of 28 research-supported hypothesized models about the relationships among sex, literacy, *APOE* ɛ4 status, and age-related cognitive decline (Table 2). We then proceeded with steps 2-7 for each of our cognitive outcomes.

**Table 2:**
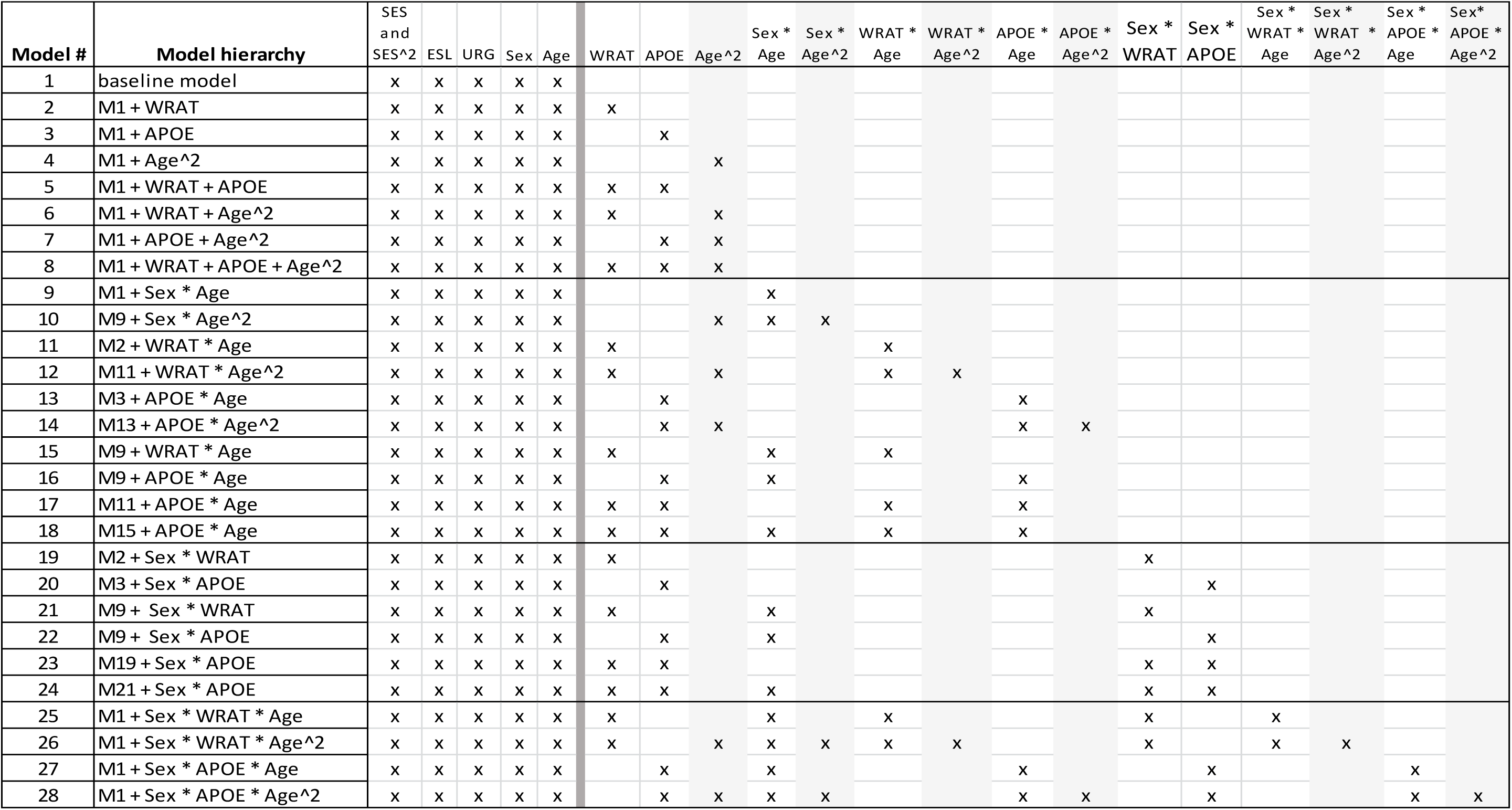
Model set fixed effects structure

| Model # | Model hierarchy | SES<br>and<br>SES^2 | ESL | URG | Sex | Age | WRAT | APOE | Age^2 | Sex *<br>Age | Sex *<br>Age^2 | WRAT *<br>Age | WRAT *<br>Age^2 | APOE *<br>Age | APOE *<br>Age^2 | Sex *<br>WRAT | Sex *<br>APOE | Sex *<br>WRAT *<br>Age | Sex *<br>WRAT *<br>Age^2 | Sex *<br>APOE *<br>Age | Sex *<br>APOE *<br>Age^2 |
| --- | --- | --- | --- | --- | --- | --- | --- | --- | --- | --- | --- | --- | --- | --- | --- | --- | --- | --- | --- | --- | --- |
| 1 | baseline model | x | x | x | x | x |  |  |  |  |  |  |  |  |  |  |  |  |  |  |  |
| 2 | M1 + WRAT | x | x | x | x | x | x |  |  |  |  |  |  |  |  |  |  |  |  |  |  |
| 3 | M1 + APOE | x | x | x | x | x |  | x |  |  |  |  |  |  |  |  |  |  |  |  |  |
| 4 | M1 + Age^2 | x | x | x | x | x |  |  | x |  |  |  |  |  |  |  |  |  |  |  |  |
| 5 | M1 + WRAT + APOE | x | x | x | x | x | x | x |  |  |  |  |  |  |  |  |  |  |  |  |  |
| 6 | M1 + WRAT + Age^2 | x | x | x | x | x | x |  | x |  |  |  |  |  |  |  |  |  |  |  |  |
| 7 | M1 + APOE + Age^2 | x | x | x | x | x |  | x | x |  |  |  |  |  |  |  |  |  |  |  |  |
| 8 | M1 + WRAT + APOE + Age^2 | x | x | x | x | x | x | x | x |  |  |  |  |  |  |  |  |  |  |  |  |
| 9 | M1 + Sex * Age | x | x | x | x | x |  |  |  | x |  |  |  |  |  |  |  |  |  |  |  |
| 10 | M9 + Sex * Age^2 | x | x | x | x | x |  |  | x | x | x |  |  |  |  |  |  |  |  |  |  |
| 11 | M2 + WRAT * Age | x | x | x | x | x | x |  |  |  |  | x |  |  |  |  |  |  |  |  |  |
| 12 | M11 + WRAT * Age^2 | x | x | x | x | x | x |  | x |  |  | x | x |  |  |  |  |  |  |  |  |
| 13 | M3 + APOE * Age | x | x | x | x | x |  | x |  |  |  |  |  | x |  |  |  |  |  |  |  |
| 14 | M13 + APOE * Age^2 | x | x | x | x | x |  | x | x |  |  |  |  | x | x |  |  |  |  |  |  |
| 15 | M9 + WRAT * Age | x | x | x | x | x | x |  |  | x |  | x |  |  |  |  |  |  |  |  |  |
| 16 | M9 + APOE * Age | x | x | x | x | x |  | x |  | x |  |  |  | x |  |  |  |  |  |  |  |
| 17 | M11 + APOE * Age | x | x | x | x | x | x | x |  |  |  | x |  | x |  |  |  |  |  |  |  |
| 18 | M15 + APOE * Age | x | x | x | x | x | x | x |  | x |  | x |  | x |  |  |  |  |  |  |  |
| 19 | M2 + Sex * WRAT | x | x | x | x | x | x |  |  |  |  |  |  |  |  | x |  |  |  |  |  |
| 20 | M3 + Sex * APOE | x | x | x | x | x |  | x |  |  |  |  |  |  |  |  | x |  |  |  |  |
| 21 | M9 + Sex * WRAT | x | x | x | x | x | x |  |  | x |  |  |  |  |  | x |  |  |  |  |  |
| 22 | M9 + Sex * APOE | x | x | x | x | x |  | x |  | x |  |  |  |  |  |  | x |  |  |  |  |
| 23 | M19 + Sex * APOE | x | x | x | x | x | x | x |  |  |  |  |  |  |  | x | x |  |  |  |  |
| 24 | M21 + Sex * APOE | x | x | x | x | x | x | x |  | x |  |  |  |  |  | x | x |  |  |  |  |
| 25 | M1 + Sex * WRAT * Age | x | x | x | x | x | x |  |  | x |  | x |  |  |  | x |  | x |  |  |  |
| 26 | M1 + Sex * WRAT * Age^2 | x | x | x | x | x | x |  | x | x | x | x | x |  |  | x |  | x | x |  |  |
| 27 | M1 + Sex * APOE * Age | x | x | x | x | x |  | x |  | x |  |  |  | x |  |  | x |  |  | x |  |
| 28 | M1 + Sex * APOE * Age^2 | x | x | x | x | x |  | x | x | x | x |  |  | x | x |  | x |  |  | x | x |

#### 2. Fit each model and check model assumptions

All models used a mixed effects structure, with the fixed effects for each model in the set as specified by Table 2, and subject specific intercepts and age-related slopes as random effects. For outcomes of AVLT Delay and BNT, logistic regression mixed models were used to address the discrete nature and ceiling/floor effects present in the data; the other outcomes used standard linear mixed effects models. For all models, SES was treated as continuous, age and SES were centered to their baseline means, and their associated quadratic terms calculated from these centered values. Each model was fit to the data by maximum likelihood, and model diagnostics were performed on the model with the most parameters and the “best fitting” model (lowest AICc value). Diagnostics included checking for homoscedastic and appropriately distributed residuals, outliers, normally distributed random effects, correlation between random effects and residuals, and overdispersion (for logistic regression). CFL and IICV were square-root transformed to address residuals issues. Stroop Color Word was removed from subsequent analysis due to several residuals violations not addressed with reasonable transforms.

Even after reasonable transformations, the following issues persisted. Correlations, between the random effects and residuals were noted for all outcomes (≤ ~0.3). Positive correlationsbetween random effects and age terms were also noted for all outcomes (≤ ~0.2), indicating the complex relationship between age and cognitive tests is not fully captured by these linear mixed effects structures. CFL also had several large positive residual outliers associated with a single subject. A sensitivity analysis for CFL removed this subject and re-performed the entire algorithm; because results did not change in any meaningful way, CFL results presented here include this subject.

#### 3. Extract model statistics

For each model in the set, the extracted model statistics included the number of model parameters (k), Akaike’s Information Criterion-corrected (AICc), and the log likelihood statistic. AICc is based on the Kullback-Leibler (K-L) divergence, which is a measure of information loss when model ‘g’ is used to approximate the true data generating model, model ‘f’ (Kenneth P. Burnham & Anderson, 2003). For data with n observations and fitted regression model ‘g’ with k parameters, the formula for AICc is (K. P. Burnham & Anderson, 2002):

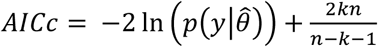

where *y* is the observed data, and 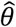 are the maximum likelihood estimates of the k parameters from model ‘g’.

#### 4. Calculate Δ_j_′s

The minimum AICc across the model set was used to calculate the difference between the best fitting model (i.e., that with the minimum AICc) and model j. For model j:
*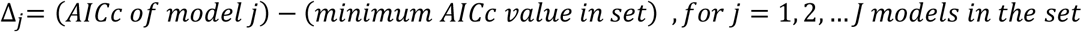*

#### 5. Calculate model weights

Δj′s for each model are used to calculate the Akaike weights (w_j_ ′s) of all models in the set. Heuristically, w_j_ represents the likelihood that model j is the K-L best model in the set. The w_j_′s helps quantify model uncertainty and are used to combine information across the set of models. w_j_ is calculated as:

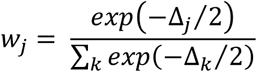

#### 6. Combine results across models

Results from each model are multiplied by their corresponding weight (w_j_), and all weighted results are then summed together for the final model averaged result. When a regression parameter does not appear in a particular model, it is set to zero in that model. Thus, for the i^th^ regression parameter in one model, the model averaged i^th^ regression parameter is:

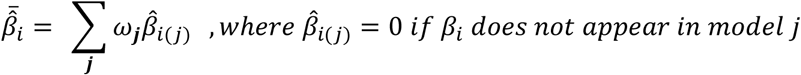

#### 7. Confidence intervals and inference using model averaged results

To facilitate multi-model inference, model-averaged estimates were evaluated using 95% CI’s obtained through non-parametric bootstrapping. For each outcome, the data used to fit the model sets was first stratified at the subject level by the total number of visits (2, 3, 4, or 5) per subject. Within each stratum, subjects were selected, with replacement, back to the number of subjects within that stratum, thus preserving the original number of subjects, observations/subject, and distribution of follow-up visits. Each bootstrap replicate went through steps 2-6; 10,000 bootstrap replicates were performed for each outcome. Bootstrap quantiles were used to calculate CI’s (using linear interpolation when necessary). Standard CI interpretation methods were used for inference about regression parameters (i.e., CI’s that did not overlap with 0 were considered significant at the α=0.05 level).

##### Bootstrap rationale

While Burnham and Anderson propose analytical methods for model-averaged inference and CI’s, these methods are predicated upon assumptions of a limiting normal distribution for the model-averaged estimate (K. P. Burnham & Anderson, 2002). Clauskens and Hjort have shown that, unless one assumes the model weights (w_j_′s) used are fixed and not random quantities, there is no guarantee of a limiting normal distribution (Claeskens & Hjort, 2008). Thus, we utilized bootstrapping methods similar to those proposed by Burnham and Anderson (K. P. Burnham & Anderson, 2002).

##### Secondary analyses Comparison of modeling methods

To illustrate how results from the above approach differ from some traditional approaches, we compared IT model averaged results for AVLT Total, AVLT Delay, and Trails B, with results obtained via single regression models determined by a best fitting model approach and by a backwards elimination approach. In both methods, the same general mixed effects model structure and use of maximum likelihood fits were used. For the best fitting approach, the model selected was that out of the candidate set which had the minimum AICc value. For backwards elimination, the starting model included all terms that appeared in any of the models within the candidate set, and the criteria for elimination was which covariate reduced AICc by the largest amount, while preserving the hierarchy of higher order terms. Elimination stopped when removal of any remaining term did not reduce AICc. For both methods, CI’s and inference from the resulting single models utilized the asymptotic normal properties of regression estimates.

*Type I error assessment*No family-wise error rate correction was performed; however, an assessment of the findings in relation to type I error rates was done. For each outcome, 27 unique non-intercept coefficients defined by the model set were examined. Thus, 270 unique coefficients were examined across all 10 outcomes: 100 main effects, 110 two-way interactions, and 60 three-way interactions. For each of these coefficient groupings, the binomial distribution was used to examine how often one would expect to detect at least the number of significant coefficients found in these analyses (at the 0.05 level), assuming the global null hypothesis that all coefficients are truly zero.

##### Software used

All analyses were performed using R version 3.4.0. Proportional hazard models were fit using ‘polr’ in the MASS package; mixed effect regression were fit using ‘lmer’ and ‘glmer’ in the lme4 package; AICc-based model statistics were calculated using ‘aictab’ in the AICcmodavg package; baseline characteristics and tests were computed using the tableone package; plots were created using the ggplot2 package; bootstrapping was performed utilizing HTCondor version 8.6.3.

## Results

### Sample Characteristics

Sample characteristics are shown in Table 1 overall and by sex. Men and women did not differ in terms of age, years of follow-up, proportion of URG or ESL, and IICV. The sexes differed on *APOE* ɛ4 count, SES, and all cognitive tests at baseline. Women performed better than men at baseline on all tests except BNT and Digit Span.

### Model-averaged Results

Model averaged coefficients and corresponding 95% CI’s for all non-intercept terms are shown in Tables 3 (AVLT Total, AVLT Delay, log_10_Trails A and log_10_Trails B, and CFL) and 4 (Digit Span forward and backward, BNT, 4-Test and 9-Test IICV). In each table, 95% CI’s that exclude 0 are identified by boldface; gray shading denotes CI’s containing 0. AICc’s and weights for all models in the set are presented in Supplemental Table 2 for all outcomes. Results are summarized below; for each outcome, results are supported by a two-panel figure (left-panel depicts 95% CIs that exclude 0 and right-panel depicts predicted outcome scores by age and selected predictors using model-averaged estimates).

**Table 3:** Model averaged parameters for all model terms AVLT Total, AVLT Delay, log_10_Trails A, log_10_Trails B, and Square-root CFL

| Model term | log10 Trails A | log10 Trails B | AVLT Total | AVLT Delayed | Sqrt CFL |
| --- | --- | --- | --- | --- | --- |
| SES | **-0.008 (-0.015, -0.002)** | **-0.013 (-0.02, -0.006)** | **0.442 (0.018, 0.86)** | **0.056 (0.001, 0.114)** | 0.047 (-0.001, 0.095) |
| SES^2 | **0.005 (2.78E-05, 0.01)** | 0.005 (-0.001, 0.012) | -0.261 (-0.537, 0.013) | -0.012 (-0.048, 0.024) | -0.003 (-0.039, 0.032) |
| URG | **0.063 (0.035, 0.091)** | **0.099 (0.067, 0.131)** | **-2.301 (-3.973, -0.677)** | -0.177 (-0.386, 0.035) | -0.003 (-0.194, 0.183) |
| Male | **0.034 (0.022, 0.053)** | **0.037 (0.021, 0.051)** | **-5.799 (-6.709, -4.908)** | **-0.64 (-0.773, -0.537)** | **-0.173 (-0.277, -0.024)** |
| Age | 2.67E-04 (-3.72E-04, 0.001) | **0.003 (0.002, 0.004)** | **-0.096 (-0.148, -0.057)** | -0.006 (-0.011, 0.001) | **0.014 (0.01, 0.017)** |
| ESL | -0.002 (-0.06, 0.054) | -0.009 (-0.064, 0.049) | -0.988 (-3.959, 1.905) | -0.176 (-0.539, 0.254) | -0.014 (-0.396, 0.379) |
| Male x Age | -1.51E-04 (-0.002, 0.001) | 1.01E-04 (-0.001, 0.002) | **-0.104 (-0.178, -0.005)** | **-0.01 (-0.021, -0.001)** | -0.001 (-0.01, 0.003) |
| one APOE ε4 | 2.69E-05 (-0.007, 0.009) | 0.001 (-0.001, 0.014) | -0.033 (-1.186, 0.052) | -0.002 (-0.134, 0.046) | 0.009 (-0.023, 0.146) |
| two APOE ε4 | -2.03E-05 (-0.028, 0.007) | 4.36E-04 (-0.008, 0.022) | -0.081 (-2.456, 0.015) | -0.014 (-0.401, 0.003) | -0.012 (-0.23, 0.043) |
| Male x one APOE ε4 | 2.41E-06 (-0.011, 0.017) | 4.39E-05 (-0.001, 0.007) | 3.41E-04 (-0.996, 0.925) | 0.003 (-0.029, 0.222) | **-0.029 (-0.368, -1.25E-06)** |
| Male x two APOE ε4 | 1.28E-04 (-0.01, 0.075) | 1.89E-05 (-0.01, 0.008) | 0.118 (-2.439, 9.44) | 0.005 (-0.368, 1.23) | -0.004 (-0.845, 0.84) |
| Age x one APOE ε4 | 2.11E-07 (-5.73E-05, 0.002) | 5.76E-08 (-1.82E-06, 9.14E-05) | -7.57E-06 (-0.004, 0.001) | -2.25E-05 (-0.006, 0.001) | 2.05E-07 (-3.72E-04, 4.92E-04) |
| Age x two APOE ε4 | -1.12E-06 (-0.003, 7.38E-06) | 1.89E-07 (-2.48E-06, 1.95E-04) | -2.79E-05 (-0.013, 0.002) | -1.42E-04 (-0.031, 0.001) | -9.33E-06 (-0.007, 7.55E-06) |
| Age^2 | 4.96E-06 (-1.48E-06, 1.32E-04) | **6.74E-05 (1.16E-06, 1.43E-04)** | **-0.001 (-0.007, -5.22E-06)** | 1.06E-06 (-5.95E-05, 3.71E-04) | -2.32E-05 (-0.001, 1.03E-06) |
| Male x Age^2 | **-8.76E-06 (-2.29E-04, -1.03E-08)** | 1.12E-05 (-5.02E-05, 1.38E-04) | 1.04E-04 (-0.004, 0.005) | -2.37E-06 (-0.001, 1.29E-05) | 2.96E-05 (-3.24E-06, 0.001) |
| Age^2 x one APOE ε4 | 3.62E-09 (-7.51E-06, 8.92E-05) | 5.41E-14 (-1.65E-20, 1.50E-07) | **-3.80E-21 (-6.02E-12, -6.18E-32)** | -6.12E-19 (-2.74E-08, 4.51E-11) | -1.66E-38 (-4.99E-27, 6.95E-28) |
| Age^2 x two APOE ε4 | 8.71E-09 (-1.30E-05, 2.01E-04) | -1.02E-13 (-2.00E-07, 2.03E-13) | -6.14E-21 (-4.74E-12, 4.94E-17) | -2.54E-18 (-2.36E-07, 3.84E-12) | -2.93E-38 (-1.90E-26, 6.40E-28) |
| Male x Age x one APOE ε4 | **-7.52E-07 (-0.005, -1.89E-11)** | **-1.01E-14 (-7.53E-07, -3.29E-28)** | 1.53E-21 (-2.83E-13, 3.13E-12) | 9.58E-18 (-1.22E-09, 6.91E-07) | 6.76E-37 (-6.22E-29, 1.24E-24) |
| Male x Age x two APOE ε4 | **2.77E-06 (3.53E-10, 0.014)** | 1.16E-14 (-4.73E-08, 1.13E-07) | 1.94E-20 (-2.38E-14, 2.35E-10) | 2.95E-17 (-2.89E-08, 1.75E-06) | 4.95E-36 (-1.74E-29, 1.79E-23) |
| WRAT reading | **0.001 (2.76E-06, 0.002)** | **-0.002 (-0.003, -0.001)** | **0.142 (0.091, 0.197)** | **0.018 (0.01, 0.025)** | **0.024 (0.018, 0.03)** |
| Male x WRAT reading | **-0.002 (-0.004, -7.27E-06)** | -0.001 (-0.003, 3.36E-05) | **0.113 (0.005, 0.213)** | 0.007 (-1.94E-04, 0.02) | **0.012 (0.001, 0.023)** |
| Age x WRAT reading | -3.70E-06 (-1.12E-04, 2.54E-05) | -6.60E-06 (-1.07E-04, 7.96E-05) | -5.13E-05 (-0.004, 0.004) | -4.21E-05 (-0.001, 2.04E-04) | 4.55E-05 (-4.26E-06, 0.001) |
| Age^2 x WRAT reading | -2.63E-08 (-6.09E-06, 5.25E-06) | 2.74E-06 (-2.96E-07, 1.61E-05) | -2.34E-06 (-2.92E-04, 2.64E-04) | -1.66E-07 (-3.77E-05, 1.99E-06) | 2.12E-06 (-5.98E-08, 6.13E-05) |
| Male x Age x WRAT reading | -1.85E-06 (-1.24E-04, 1.03E-04) | 6.46E-07 (-1.31E-04, 1.31E-04) | 0.001 (-0.003, 0.009) | 6.11E-05 (-5.62E-05, 0.001) | -3.18E-05 (-0.001, 1.03E-04) |
| Male x Age^2 x WRAT reading | 2.27E-08 (-7.72E-06, 8.58E-06) | -3.08E-06 (-2.16E-05, 1.22E-07) | -2.70E-05 (-0.001, 3.34E-04) | 1.28E-07 (-6.11E-06, 3.61E-05) | -2.32E-06 (-7.99E-05, 1.35E-06) |
| Male x Age^2 x one APOE ε4 | -1.06E-08 (-2.30E-04, 6.12E-07) | -1.10E-16 (-6.63E-09, 1.58E-10) | 3.71E-22 (-8.99E-16, 1.21E-12) | -3.42E-19 (-7.80E-08, 2.43E-11) | **-1.19E-37 (-5.49E-25, -1.24E-49)** |
| Male x Age^2 x two APOE ε4 | -4.06E-08 (-0.001, 3.08E-05) | 1.26E-15 (-9.45E-13, 8.32E-08) | -2.84E-21 (-4.52E-11, 6.42E-20) | -6.96E-18 (-4.33E-06, 2.47E-13) | 2.66E-37 (-2.49E-28, 3.36E-25) |

Note: "E" represents "x10<sup>^</sup>"; e.g. 1.23E-04 = 1.23x10<sup>-4</sup>
95% CI's that do not overlap 0 (are significant) are identified by bold-face font; gray shading denotes CIs that overlap 0.

#### Memory

For AVLT Total, Model 21 was best-fitting, contributing a weight of .562 to model averaging; Model 26 contributed .183 and Model 25 contributed .139 (weights in supplemental Table 2); all other model weights were under .05. Significant interactions included ɛ4 count 1*quadratic age, sex*age, and sex*literacy (left-panel, Figure 1). Figure 1 (right-hand panel) depicts AVLT Total model-averaged predicted performance for the latter two interactions since the estimated beta for the interaction with age and ɛ4 count 1 was ostensibly 0. There’s a larger gap in AVLT Total scores for high vs low literacy in men than in women and men show faster AVLT Total age-related decline than women.

**Fig. 1.**
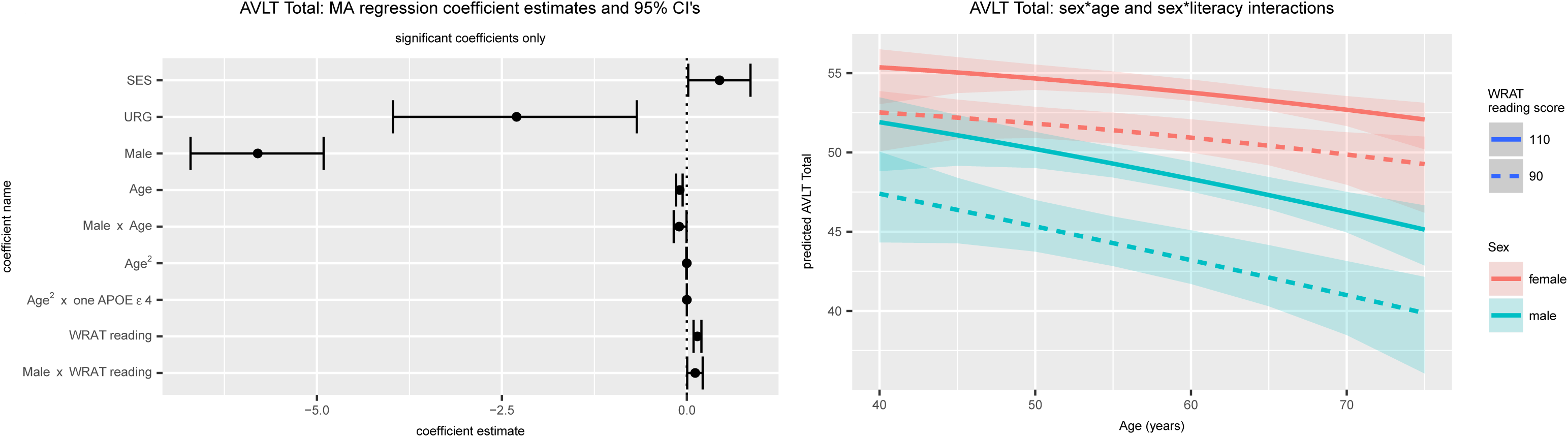

The best fitting model for AVLT Delay was also Model 21 (weight=.531), followed by Model 15 (.123) and Model 25 (.119). Variability in AVLT Delay was explained largely by SES, sex, literacy and age, with age-related AVLT Delay decline steeper for men than women (see Supp. Fig. 1A for CI’s and predicted trajectories).

#### Executive function/ Working Memory

The best fitting models for Trails B were Models 6 (weight=330), 26 (.300), 12 (.124), and 19 (.109). Variability in Trails B was explained by SES, URG status, sex, literacy, age (linear and quadratic), and a 3-way interaction with sex* *APOE* ɛ4 count 1*age (left-hand panel, Figure 2A). Though significant, the estimated beta for the sex*age* *APOE* interaction was again close to 0. Predicted age-related trajectories show slower times for men than women and lower vs higher literacy (right-hand panel, Figure 2A; no predicted values are shown for men and *APOE* ɛ4 count=2 due to the small cell size (n=4)).

The best fitting models for Digit Span Backward were Models 13 (weight=.583), 15(.168), 20(.117), and 15(.104). Variability in Digit Span Backward was explained by quadratic age, literacy, and sex with greater sex differences at higher literacy levels (Supp. Figure 1B).

**Fig. 2a.**
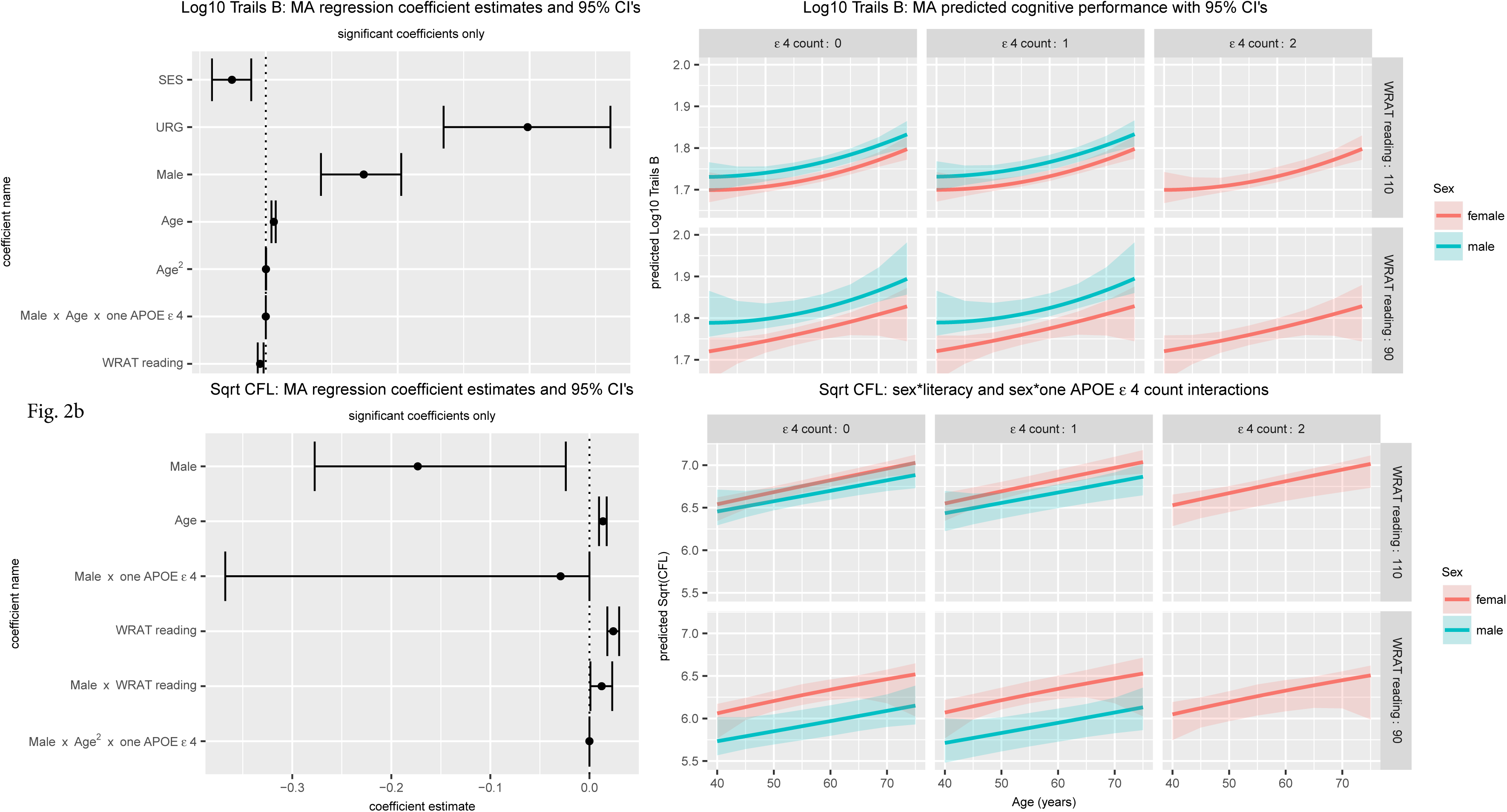

#### Attention

The best fitting models for Trails A were Models 19 (weight=.574), 21 (.248), and 25 (.064). Supplemental Figure 1C depicts significant beta estimates and their CI’s (left panel) and predicted values vs age, stratifying by sex and literacy (high vs low; right panel). Predicted values indicate that women of high literacy perform worse on Trails A then women of low literacy while the opposite is true for men. Although the interactions of sex*quadratic age, and sex*age* *APOE* ɛ4 counts of 1 or 2 were statistically significant, all had estimated betas extremely close to 0 (Table 4; Supp. Fig. 1C).

**Table 4:**
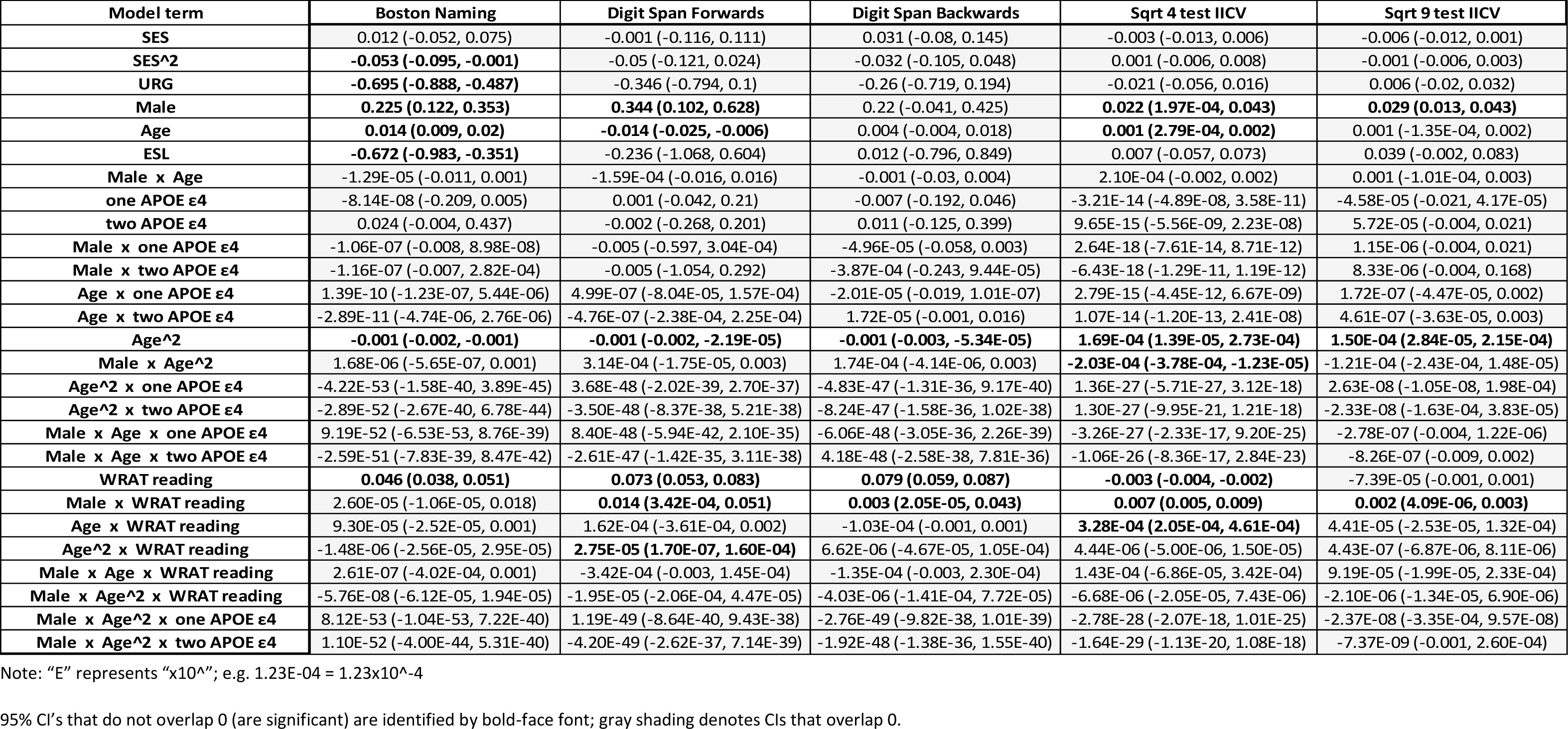
Model averaged parameters and 95% CI’s for all model terms and CFL, BNT, Digit Span Forwards, Digit Span Backwards, 4 Test IICV and 9 Test IICV

The best fitting models for Digit Span Forward were Models 6 (weight=.307), 26 (.234), 19 (.163), and 12 (.124). Variability in Digit Span Forward was explained by sex, age (linear and quadratic) and literacy; significant interactions included sex*literacy and literacy*quadratic age. The latter beta estimate was near zero; sex differences in the outcome are negligible at low literacy levels while men outperform women at high literacy levels (Supp. Figure 1D).

#### Language

The best fitting models for CFL were Models 13 (weight=.416) and 14 (.251). Variability in CFL was explained by age, sex, literacy, and interactions sex* *APOE* count 1, sex*literacy, and sex* *APOE* count 1*quadratic age, although the three-way beta estimate was essentially 0 (Figure 2B). Women did better overall and sex differences were smaller in ɛ4 count=0 than count=1. All improved with age. The best fitting models for BNT were Models 6 (weight=.630), 12 (.239), and 8 (.128). Variability in BNT was explained by SES, URG status, sex, age, ESL status, and literacy level with higher literacy, SES, male sex and older age associated with better performance, while URG and ESL status were associated with lower BNT scores (Supp. Figure 2).

#### Intraindividual variability

The best fitting model for 4-Test IICV was Model 26 (weight=0.959); all other models contributed <0.05 to the parameter weights. Variability in 4-Test IICV was accounted for by sex, age, and literacy, with significant sex*quadratic age, sex*literacy, and literacy*age interactions (Figure 3). At lower levels of literacy, women show higher 4-Test IICV than men and IICV declines with age; in contrast, IICV increases with age at higher literacy levels and is at times higher among men than women (Figure 3). The best fitting model for 9-Test IICV was also Model 26 (weight=0.890); significant parameters indicate that 9-Test IICV patterns differed most between men with high and low literacy levels (Supp. Figure 3).

**Fig. 3.**
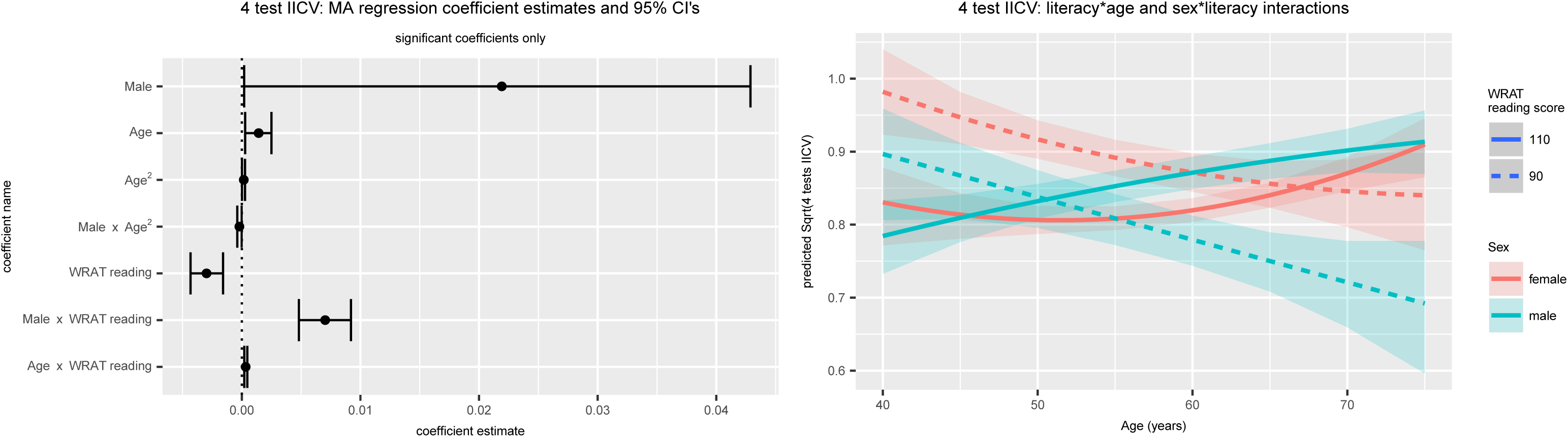

#### Model comparisons

In secondary analyses of 3 outcomes (AVLT Total, AVLT Delay, and Trails B), we compared estimated betas and 95% CI’s between the IT, best fit, and backward selection approaches for coefficients that were significant in any of the methods, per outcome. Results differed most across model selection approaches for Trails B, with 6 of 11 terms inconsistently significant across methods (see Figure 4 for point estimates and CI’s for the three approaches for Trails B and Supplementary Figure 4 for AVLT Total and Delay).

**Fig. 4.**
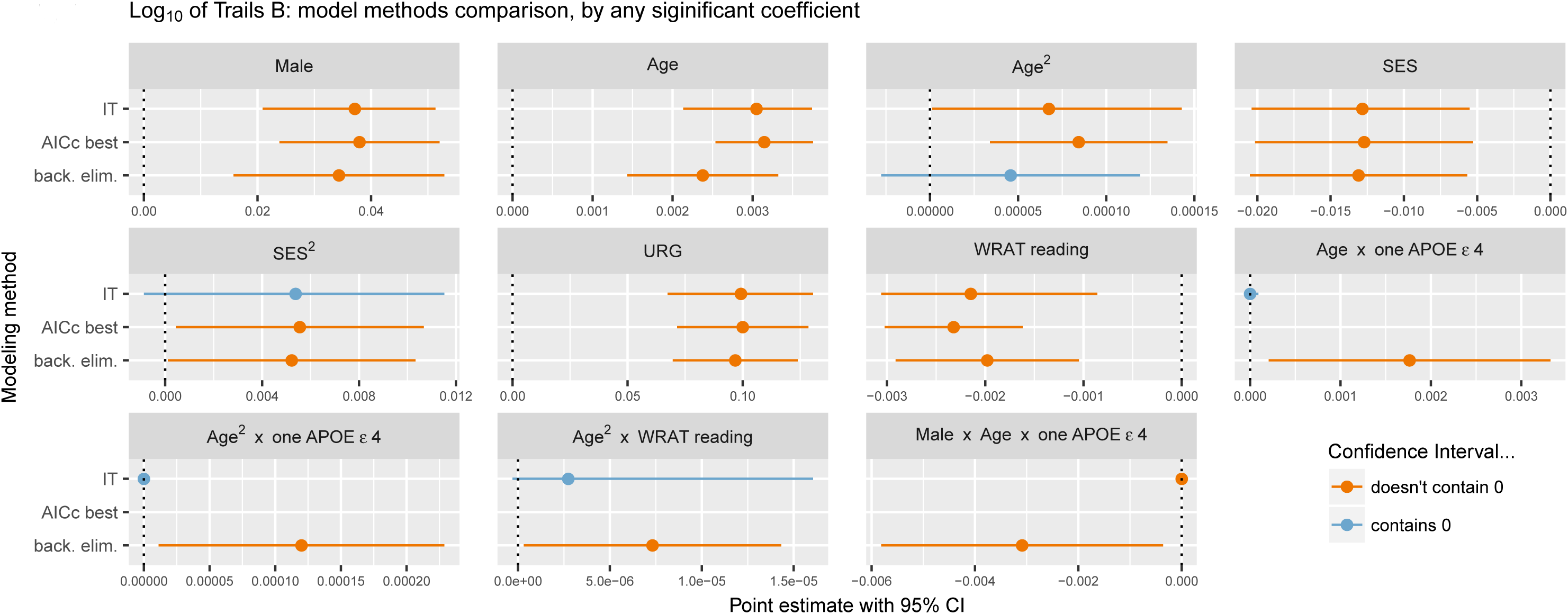

#### Type I error assessment

Forty-two percent of the 100 unique main effects examined across all 10 outcomes were significant at the 0.05 level. Under the global null hypothesis for all coefficients, the probability of detecting at least 42 coefficients is 2.50x10-^29^. Fifteen (13.6%) of 110 unique two-way interactions were significant, corresponding to a probability of 0.00012 under the global null. Four (6.67%) of 60 unique three-way interactions were significant, corresponding to a 0.180 probability under the global null.

## Discussion

In this study, we used information-theoretic (IT) model averaging techniques to characterize how sex, *APOE* ɛ4 carrier status, and literacy modify age-related cognitive and IICV trajectories in a sample that was middle-aged and free of clinical impairment at baseline assessment (mean(sd) age at baseline and last visit: 53.7(6.6) and 62.8(6.8), respectively). We observed age-related declines for all cognitive outcomes except the two language-related measures (CFL and BNT). Age-related declines in IICV were associated with lower literacy levels while IICV tended to increase with age among participants with higher literacy. Significant but small quadratic age effects were observed for a few outcomes. *APOE* ɛ4 count showed significant but small modifying effects on age-related trajectories on four outcomes. Sex and literacy were consistently significant predictors of measures of memory, executive function, working memory, language and intra-individual cognitive variability including significant main effects, and sex*age and/or sex*literacy interactions for a subset of outcomes.

Compared to those with no *APOE* ɛ4 alleles, carriage of one or two ɛ4 alleles is associated with greater risk of AD (Neu et al., 2017) and faster or earlier cognitive decline in certain domains. For example, in a sample of cognitively normal adults (mean baseline age ~60 years, followed an average of ~5 years), Caselli et al (Caselli et al., 2009) reported accelerated age-related decline on AVLT Delay among *APOE* ɛ4 carriers (vs non-carriers) beginning prior to age 60. Predicted annual rate of AVLT Delay change, however, was very small for carriers and non-carriers in age-ranges similar to our sample (e.g., 50-59, 0.07 vs 0.08 and 60-69, 0.04 vs −0.03 for non-carriers vs carriers, respectively). In a meta-analyses, Wisdom and colleagues (Wisdom et al., 2011) also reported significant yet small differences in age-related decline among *APOE* ɛ4 carriers relative to non-carriers (Cohen’s d estimated effect sizes <.20; (Wisdom et al., 2011)). Results in our sample also showed small effects. Specifically, significant age-modifying effects of one *APOE* ɛ4 allele (vs 0) were noted for AVLT Total, Trails A and B, and CFL, although beta estimates for these interactions were essentially zero and not clinically meaningful.

Previous research has also shown sex differences in risk of MCI (Roberts et al., 2012) and AD (e.g., (Altmann et al., 2014)), and rates of decline (e.g., (McCarrey, An, Kitner-Triolo, Ferrucci, & Resnick, 2016; Mortensen & Høgh, 2001)), with other studies showing evidence of potential sex*ɛ4 interactions (Payami et al., 1996). Sex*age interactions showed faster decline in men for both AVLT measures. These results are consistent with those reported for the Mayo Clinic Study of Aging in which memory, memory decline, and hippocampal volume were worse among men compared to women (Jack CR, Jr, Wiste HJ, Weigand SD, & et al, 2015) A small number of significant sex* *APOE \**age interactions were observed, but beta estimates were essentially 0. As we follow the WRAP cohort into their 70’s, larger effects may emerge for these complex interactions.

Literacy level as measured by word reading tasks is considered a proxy for verbal intellectual ability and quality of education; lower literacy has been associated with faster memory decline (Manly et al., 2003). Higher literacy levels have also been shown to be associated with resilience to *APOE* ɛ4 - related cognitive decline (Kaup et al., 2015; Vemuri P, Lesnick TG, Przybelski SA, & et al., 2014). In our sample, however, literacy did not modify effects of *APOE* ɛ4 or ageon cognition. The benefits of high literacy were stronger for men than women for AVLT Total, Trails A, CFL and Digit Span.

IICV has been shown to vary across the lifespan (Salthouse & Soubelet, 2014). In our data, lower literacy was associated with steady age-related decline in 4-Test IICV with higher IICV consistently observed among women; at higher literacy levels, IICV was generally higher among men and increased with age for both sexes. Our 4-Test IICV is the same as used in previous WRAP analysis and similar to what has been used in other analyses (Holtzer et al., 2008; Anderson et al., 2016, Gleason et al, 2017) which indicate higher IICV predicts MCI and AD. As IICV is developed further as a potential cognitive marker for risk of later decline in the AD continuum, future studies should examine whether risk-indicating value is constant across underlying demographics such as age, sex, and literacy.

Traditional model selection methods such as stepwise regression are prone to overfitting the data, producing overconfident estimates with standard errors that do not account for the degrees of freedom in the search process (Hastie, Trevor, Tibshirani, Robert, & Friedman, Jerome, 2009). Shrinkage methods, such as Lasso, can help select important predictors with respect to the outcome, but parsimonious models and predictive accuracy are the typical goal, and statistical inference can be difficult (Hastie, Tibshirani, & Wainwright, 2015). Bayesian model averaging combines information from posterior distributions of parameters of interest across several models, weighting each by its posterior model probability (Hoeting, Madigan, Raftery, & Volinsky, 1999). However, it is important in Bayesian methods to formulate reasonable prior distributions for all parameters and model probabilities, and this can be prohibitive when the set of models under consideration is large (Claeskens & Hjort, 2008). By using the IT approach in this paper, we obtained the robustness benefits of model averaging without the overhead of Bayesian methods, while still yielding familiar statistical outputs that support inferences (i.e. point estimates, CI’s). IT methods also allow examination of several scientific questions of interest; when only one or a few model structures are necessary to handle the scientific question(s) of interest, IT methods may be of little value considering the extra complexity and time required. However, since age-related decline is influenced by many factors, the IT approach offers an attractive framework for evaluating competing hypotheses about modifiers of decline.

Our secondary analyses suggest that the IT approach may guard against overidentifying and overestimating effects compared to traditional methods. First, in our comparison of the IT approach with best fit and backwards elimination approaches, main effects estimates between the three methods were generally very similar, though the IT method tended to have the widest confidence intervals. Relationships between the three methods were more complex for quadratic effects and interactions. Backwards elimination commonly “found” interactions that best fit did not, while IT tended to attenuate the estimated interaction coefficient ostensibly to zero. Second, the high numbers of significant two-way interaction effects detected in the IT method were well above those expected from random chance under the global null, lending confidence to conclusions about significant IT model-averaged effects. For all 4 significant three-way interactions, the IT method estimates have very tight CI’s that are extremely close to zero, especially compared to estimates that were found by backwards elimination. This suggests that IT methods might further guard against overconfident results by estimating some significant coefficients to be ostensibly zero.

The generalizability of our results is limited by cohort characteristics, including that our sample is relatively young, highly educated, enriched for AD risk, has limited follow-up on participants from URG, and has few males homozygous for ɛ4. In addition, use of *APOE* ɛ4 count is just one of many possible ways of parameterizing *APOE*-associated risk.

## Conclusions

While carriage of an *APOE* ɛ4 allele is known to be the greatest modifier of age-related risk for late onset AD, many factors may interact to hasten or slow the rate of cognitive decline during mid-life. When several predictors and their interactions are considered simultaneously, model selection procedures can result in overestimation of significance of certain parameters and loss of information about others that are accounting for similar amounts of variability. The IT model averaging approach offers a framework that allows results from multiple plausible hypotheses to provide weighted model-averaged parameter estimates and CI’s, with CI’s excluding zero interpreted as significant. The application of this methodology to the WRAP sample suggests that age-related trajectories are modified more by sex and literacy levels than by *APOE* ɛ4 allele count in this age range. Future applications of the IT methodology will examine the interplay of sex and literacy with other potential cognitive trajectory-modifying variables such as polygenic risk, AD biomarkers, or lifestyle factors (e.g., exercise or diet).

## Acknowledgments

This research was supported by the National Institutes of Health awards R01 AG027161, R01 AG054059, UL1 TR000427 and by donor funds including the Wisconsin Alzheimer’s Institute Lou Holland Fund. Portions of this research were supported by resources at the Wisconsin Alzheimer’s Institute, the Wisconsin Alzheimer’s Disease Research Center and the Geriatric Research Education and Clinical Center of the William S. Middleton Memorial Veterans Hospital, Madison, WI. Any opinions, findings, and conclusions or recommendations expressed in this material are those of the authors(s) and do not necessarily reflect the views of the NIH or the Veterans Administration. We gratefully acknowledge the WRAP study team who have carefully acquired the longitudinal data, and the WRAP participants who make this research possible.

